# Single-stranded HDR templates with truncated Cas12a binding sequences improve knock-in efficiencies in primary human T cells

**DOI:** 10.1101/2024.09.11.608426

**Authors:** Ana-Maria Nitulescu, Weijie Du, Viktor Glaser, Jonas Kath, Robert Greensmith, Nanna Steengaard Mikkelsen, Maik Stein, Rasmus O. Bak, Michael Kaminski, Dimitrios L. Wagner

## Abstract

Non-viral gene editing via CRISPR-Cas12a offers an alternative to Cas9-based methods, providing better targeting of AT-rich regions, simplified guide RNA manufacturing, and high specificity. However, the efficacy of editing outcomes is subject to various factors, with template format playing a crucial role. Currently, the predominant non-viral template format for inducing homology-directed repair (HDR) after nuclease-induced DNA breaks is double-stranded DNA (dsDNA), which is toxic when transfected at high doses. Previous studies have demonstrated that using single-stranded DNA (ssDNA) with flanking double-stranded Cas-target-sequences (CTS) as a repair template for Cas9-mediated gene editing can mitigate this toxicity and increase knock-in efficiency. Here, we investigate CTS design for AsCas12a Ultra by exploring PAM orientation and binding requirements of the Cas12a-crRNA complex. Additionally, we rule out *in vitro* ssDNase activity of AsCas12a Ultra under cell-physiological Mg^2+^ conditions. Finally, we showcase the advantage of using ssDNA with double-stranded CTS end modifications (ssCTS) at high doses for delivering clinically relevant transgenes of varying sizes into three T-cell receptor-CD3 complex genes (*TRAC*, *CD3*ζ*, CD3*ε), achieving up to 90% knock-in rates for a 0.8kb insert at the *CD3*ε locus. Overall, AsCas12a Ultra and ssCTS donors represent a platform for highly efficient knock-in in primary human T cells with minimal toxicity.

## 2. Introduction

Adoptive transfer of engineered T cells expressing synthetic antigen receptors, such as chimeric-antigen receptors (CARs), is an effective approach for second- or third-line treatment for B-cell malignancies (1, 2). Currently, all approved CAR-T cell products are manufactured based on nontargeted integration of transgenes using retro- or lentiviral vectors. However, this process is associated with significant expenses linked to the production and testing of clinical grade viral vectors (3). To reduce these costs, clinical trials have commenced using CAR-T cells produced with non-viral transposase systems. Like retroviruses, transposases integrate their cargo semi-randomly into the chromosomes, and in the case of a hyperactive version of the transposase piggyBac, this has contributed to the development of CAR-positive T cell malignancies (4).

Reprogramming of cells using the CRISPR-Cas system is an attractive non-viral gene transfer alternative, enabling precise, targeted genomic modifications that enhance the quality and safety of the cellular product by reducing the risk of insertional mutagenesis (4, 5). Furthermore, while randomly integrating vectors depend on exogenous promoter-driven transgene expression, non-viral gene editing can harness endogenous gene regulation to enhance product potency. For instance, knock-in into the *T-cell Receptor (TCR) Alpha Constant* (*TRAC*) locus has been shown to improve CAR-T cell potency and persistence in preclinical mouse models of B-cell acute lymphoblastic leukemia (B-ALL) (6). Ongoing clinical studies are investigating the potency of *TRAC*-replaced CAR-T cells in treatment-refractory large B-cell lymphoma (7). Gene editing of other TCR/CD3 complex genes, such as *CD3*ζ and *CD3*ε, has also been proposed to create potent CAR-T cells (8, 9) as well as TCR fusion constructs (TRuC) (10–12). We previously demonstrated that *CD3*ε gene editing with TRuC is a strategy uniquely suited for redirection of immunosuppressive immune cells, called regulatory T cells (Tregs) (13). Reprogramming Tregs to recognize allo-antigens, such as HLA-A2, holds significant potential to suppress allo-mediated rejection in solid organ transplantation (14) and to reduce or replace hazardous long-term immunosuppression in patients. Consequently, nonviral gene editing is a promising gene transfer modality to manufacture redirected T cell products with enhanced fitness for diverse medical applications.

The efficacy of site-specific gene transfer using CRISPR-Cas gene editing depends on various factors, such as the targeted locus, the specific programmable nuclease, guide RNA (gRNA) selection, and the homology-directed repair template (HDRT) (15, 16). The most commonly used non-viral template formats include plasmids (17–21), linear double-stranded DNA (dsDNA) (9, 22–28), and single-stranded DNA (ssDNA) (22, 29). Previous studies have shown that electroporation of cells with ssDNA exhibits reduced cell toxicity compared to dsDNA templates (22). However, aside from the use of small single-stranded oligonucleo-tides (ssODNs) for point mutation repair and smaller inserts (30), knock-in rates using ssDNA for larger transgenes are generally low. To boost CRISPR-Cas9 editing efficacy with ssDNA, Shy et al. proposed to integrate truncated Cas-target-sequences (tCTS) into the ends of ssDNA HDRTs (31). These sequences are intended to enhance template delivery into the nucleus by serving as binding sites for the Cas9 protein, which contains nuclear localization signals (NLS) (23). The truncated format of the CTS is designed to prevent Cas-mediated cleavage of the template. To date, ssDNA with double-stranded tCTS (ssCTS) has not been adapted from the Cas9 system to other programmable nucleases.

In addition to considering the HDRT format, careful selection of the nuclease is warranted to ensure compatibility and efficient DNA double-stranded break (DSB) induction within the target region. Given that most mammalian genes are GC-rich, the presence of the requisite 5’-NGG-3’ protospacer adjacent motif (PAM) enables targeting *Streptococcus pyogenes* Cas9 (SpCas9) to specific locations. This, coupled with the high efficacy and extensive characterization of the enzyme (32–34), has established SpCas9 as the preferred choice for genome editing. However, alternative nucleases like *Acidaminococcus species* Cas12a (AsCas12a) present advantageous properties over SpCas9 for specific applications (35). Unlike SpCas9, AsCas12a creates staggered-end DSBs distal from a T-rich PAM, which helps guide the precise alignment of complementary DNA sequences during HDR (35, 36). Moreover, the nuclease operates effectively with only a short single crRNA of 42 nucleotides, combining the roles of both crRNA and tracrRNA, which is easier to manufacture via solid state synthesis than gRNAs of 100-nucleotide length required for SpCas9 (37). Furthermore, studies have shown that AsCas12a is less tolerant for mismatches than SpCas9, thereby reducing unintended off-target effects (37, 38). Despite its advantages, the widespread adoption of Cas12a as a genome editing tool has been hindered by its comparatively lower editing efficiency in living cells (39). To address this limitation, Zhang et al. developed an enhanced version of Cas12a, known as AsCas12a Ultra, by introducing two-point mutations, M537R and F870L to boost its activity (40). Their study demonstrated that these mutations significantly improved efficacy while maintaining the high intrinsic dsDNA-specificity of the nuclease. Previous studies demonstrated that both LbCas12a and AsCas12a enzymes can display indiscriminative ssDNase function *in vitro* – a function that is activated following successful cleavage of the dsDNA target (41, 42). This feature has sparked the development of *in vitro* diagnostic assays (43), however, it could be detrimental for genome editing efficiency with ssDNA HDRTs due to donor degradation *in cellulo*. It is unclear whether AsCas12a Ultra shares this feature of other Cas12a enzymes. Consequently, AsCas12a Ultra offers a putative alternative to SpCas9 that has the potential to expand the therapeutic genome editing landscape, although the ssDNase activity could represent a caveat for its use with ssDNA HDRTs. In this study, we demonstrate highly efficient virus-free gene editing of T cells using modified ssDNA templates and AsCas12a Ultra. First, we examined various AsCas12a binding motifs as end modifications in dsDNA templates. After optimization of dsDNA templates, we incorporated different promising CTS-motifs into ssDNA HDRTs and performed investigations regarding a previously described Mg^2+^-dependent ssDNase activity of the AsCas12a (41) *in vitro*. Digestion assays demonstrated that neither intact nor truncated CTS-modified ssDNA templates were digested under physiological Mg^2+^ concentrations. Finally, we assessed the HDR efficacy and toxicity of different non-viral DNA templates (dsDNA or ssDNA with or without CTS) encoding clinically relevant transgenes for cancer, autoimmune disorders, and transplantation medicine. Regardless of the transgene or the specific locus, using ssCTS and AsCas12a exhibited improved HDR efficiencies by 3-10 fold over unmodified ssDNA and sustained high cell viability even at the highest template concentration employed.

## 3. Materials and Methods

### PBMC isolation and T cell enrichment

The research was conducted in accordance with the Declaration of Helsinki. Peripheral blood samples were collected from consenting healthy adult individuals under the approval of the Charité ethics committee (approval code EA1/052/22). Peripheral blood mononuclear cells (PBMCs) were isolated through density-gradient centrifugation. For this, 50-mL LeucoSEP™ tubes (Greiner Bio-One GmbH) were used, to which 15 mL of BioColl® separating solution (Ficoll) (Bio&SELL GmbH) à tube were added. The tubes were then quickly spun down, in order for the solution to pass the porous filter. Fresh heparinized whole blood was mixed in a 1:1 ratio with sterile phosphate-buffered saline (PBS) (Gibco) and poured into the Ficoll-containing tubes. Centrifugation was carried out at 1000 x g for 20 minutes employing minimal break speed (acceleration 6 and deceleration 3). The mononuclear cell layer was then collected, diluted in sterile PBS, and subjected to two centrifugation steps with break at 300 x g for 10 minutes, each with subsequent removal of the supernatant. Afterwards, the PBMC pellet was resuspended in 50 mL of PBS and counted using the CASY cell counter (OMNI Life Science GmbH & Co. KG). Finally, PBMCs were positively enriched for CD3+ T cells using magnetic column enrichment with human CD3 microbeads, following the manufacturer’s recommendations (LS columns, Miltenyi Biotec, Germany).

### Cell culture

T cells were cultured in 44.5% Click’ medium (Irvine Scientific), 44.5% Advanced RPMI medium 1640 (Gibco), including 10% heat-inactivated fetal calf serum (FCS) (Sigma-Aldrich), 1% GlutaMAX (100X) (Gibco), recombinant IL-7 (10 ng/mL) and IL-15 (5 ng/mL) (CellGenix GmbH). T cell activation was carried out for 48 hours on tissue culture plates coated with anti-CD3/28 antibodies. For this, 24-well-tissue-culture plates (Corning) were incubated overnight at 4°C with 500 µL/well of sterile ddH2O supplemented with 1 μg/mL anti-CD3 monoclonal antibody (mAb) (clone OKT3; Invitrogen) and 1 μg/mL anti-CD28 mAb (clone CD28.2; BioLegend). Afterward, the plates were rinsed twice in PBS and once in RPMI without allowing the wells to dry. T cells were then seeded at a density of 1-1.5 x 10^6^ cells per well and cultured at 37°C, 5% CO_2_.

### Design of plasmids encoding homology-directed repair templates

The majority of DNA templates used in this study were generated from previously published plasmids (9, 13, 27). These encoded for receptors or fusion constructs, such as CD19-CARs and HLA-A2-TRuC, intended for integration into specific loci of primary human T cells, including the *T-cell receptor* α *constant chain* (*TRAC*), *CD3*ζ (*zeta chain of the CD3 complex*), or *CD3*ε (*epsilon chain of the CD3 complex*). Both the CAR and TRuC constructs were flanked by approximately 400-bp-long HAs complementary to the genomic DNA sequence next to the cutting site of interest as previously described (9, 13, 22, 27). For the *TRAC*-directed CAR templates, a second-generation CAR design was employed. This included a CD19-binding single-chain variable fragment (scFv) with an IgG1 hinge region followed by a CD28 transmembrane and co-stimulatory region and a CD3ζ domain. The heavy chain of the scFv was connected to the light chain via a triple glycine and 4 serine-rich (3xG4S) linker. In the case of the *CD3*ζ-directed CARs, no exogenous stimulatory CD3ζ was integrated into the design, as the endogenous CD3ζ is recruited for CAR assembly (9). All CAR constructs featured a porcine teschovirus-1 2A (P2A) sequence positioned upstream of the scFv domain, and following this, a membrane leader sequence. The *TRAC* construct also included a bovine growth hormone-derived polyadenylation site (bGH poly A). No poly A was added to the *CD3*ζ-directed templates because the CAR transgene was inserted in-frame into the gene encoding the CD3ζ protein (9). Of the two *CD3*ζ-directed CARs used, one contained a Myctag. In contrast to the CARs, the TRuC template was comprised solely of a Myc-tag, an HLA-A2-binding scFv, and two 3xG4S linkers – one for linking the light and heavy chains of the antibody fragments, and one for binding the endogenous CD3ε chain. All plasmid sequences are listed in Supplementary Table 1, the original *CD3*ζ*-*HDRT and the original *TRAC-*HDRT are available via Addgene (CD3ζ -truncCARgsg: Addgene ID 215759, TRAC-Cas12a: 215769).

### In-Fusion cloning strategy of plasmids

The cloning of plasmids encoding HDRTs was conducted using the two-fragment In-Fusion method following the manufacturer’s protocol (Clontech, Takara Bio). For the *CD3*ζ-directed CD19-CAR construct containing a Myc-tag, an In-Fusion cloning strategy was planned with SnapGene (from Insightful Science; snapgene.com). The CAR transgene and the backbone were PCR amplified from previously published plasmids (9, 13), and the resulting products were purified using the DNA Clean and Concentrator-5 Kit following the manufacturer’s instructions (Zymo Research). In-Fusion reactions were prepared in 5 µL volumes at a 1:3 vector to insert molar ratio. Subsequently, 2.5 µL of the In-Fusion reaction mixture was transformed into Stellar Competent E. coli (Takara Bio) in 10 µL reactions and then plated on LB (Carl Roth GmbH) broth agar plates supplemented with ampicillin (Sigma-Aldrich). Following colony PCR for size validation, preferred clones were cultured overnight at 37°C, 200 rpm in 3–5 mL ampicillin-containing bacterial cultures. Plasmids were purified using the ZymoPURE Plasmid Mini Prep Kit (Zymo Research). Lastly, sequence confirmation of HDR donor template-containing plasmids was accomplished through Sanger Sequencing (LGC Genomics, Berlin).

### Primer and oligo design for CTS HDR templates

Primers for generating HDR templates with various CTS motifs were designed as recently described (23, 31) and synthesized by IDT. In the case of dsCTS, all primers were comprised of a 16 bp-long buffer sequence, the specific gRNA or crRNA target sequence with mismatches ranging from 0 to 12 bp, a PAM, and the complementary sequence for amplification. A similar design was employed for generating ssCTS, except for the use of either a shorter 4 bp-long buffer sequence or no buffer sequence at all. Moreover, primers for ssDNA production were either 5’ or 3’ phosphorylated. For the generation of ssCTS templates with double-stranded ends, complementary oligos were designed and synthesized by IDT. All oligos included the specific guide RNA target sequence with 0, 2, 4 or 6 bp mismatch, the PAM and the complementary sequence to the homology arms. Only some oligos contained a buffer sequence. All primer and oligo sequences are listed in supplementary table 2.

### Generation of double-stranded and single-stranded DNA for homology-directed insertion of receptors

The HDR templates were amplified from plasmids by PCR using the KAPA HiFi HotStart 2x Readymix (Roche) with reaction volumes of either 500 μL or 1000 μL. The resulting dsDNA amplicons were concentrated and purified using paramagnetic beads (AMPure XP, Beckman Coulter Genomics). In this process, PCR products were mixed with the beads in a 1:1 ratio, incubated at room temperature for 10 minutes, and then placed in a DynaMag^TM^-2 stand (Invitrogen, Thermo Fisher Scientific) for another 10 minutes. The bead-nucleic acid mix was washed 2 times under sterile conditions with freshly prepared 70% ethanol while still on the magnet. Finally, the DNA was eluted in 3 μL of nuclease-free water per 100 μL PCR product. The concentration of the nucleic acid was determined either using the Nanodrop ND-1000 spectrophotometer (Thermo Fisher Scientific) or the Qubit 2.0 fluorometer (Invitrogen, Life Technologies). For the generation of ssDNA templates via enzymatic digestion, the “Guide-it Long ssDNA Production System v2” kit was used according to the manufacturer’s instructions (Takara Bio). In this process, the PCR products amplified with one phosphorylated primer were used. After ssDNA production, templates were purified and concentrated similar to the dsDNA cleanup, using AMPure XP beads. For the generation of double-stranded CTS ends on the ssDNA, complementary oligos were added at a 6:1 molar ratio of oligos to nucleic acid. For the annealing, the mixed solutions were incubated at 94°C for 2 minutes, followed by 70°C for 1 minute, and finally room temperature for 15 minutes. The concentration of all DNA templates was adjusted according to the knock-in experimental setup planned (range of 0.5-4 μg/μL).

### Ribonucleoprotein formulation and mix with DNA templates

Prior to the electroporation RNPs were formed by mixture and incubation of 0.5 μL Poly-Lglutamic acid (PGA) (Sigma-Aldrich, 100 mg/mL), 0.48 μL of crRNA or sgRNA (synthesized by IDT, 100 μM) and 0.4 μL of Alt-R® AsCas12a (Cpf1) Ultra (synthesized by IDT, 64 μM) or Alt-R S.p. Cas9 Nuclease V3 (synthesized by IDT, 61 μM) for 15 minutes at room temperature. After the incubation, the RNPs were placed on ice until further use. For the electroporation of T cells with HDRTs containing CTS motifs, the DNA was pre-incubated for 5-10 minutes with the RNPs. Depending on the experimental setup, different HDRT amounts were used as indicated. Information regarding the crRNA or sgRNA target sequences can be found in supplementary table 3.

### Electroporation

Non-viral knock-ins in T cells were performed as recently described (27). 48 hour-stimulated primary T cells were harvested, counted and washed twice in sterile PBS for 10 minutes, first at 150 x g and then at 100 x g. Afterwards, 1 x 10^6^ cells were resuspended in 20 μL P3-buffer (Lonza) per electroporation reaction and added to the RNP-HDRT mix. The suspension was carefully transferred to a 16-strip electroporation cuvette, which was then tapped on the bench repeatedly to guarantee proper positioning of fluids within the strip and the absence of bubbles that would interfere with the electric current. The cells were electroporated with the 4D-Nucleofector device (Lonza) using the program EH-115 according to previous reports (22, 27). Quickly after, 90 μL of pre-warmed medium were added per electroporation reaction, and the strip was incubated for 10 minutes at 37°C. Lastly, the cells were seeded to a 96-well round-bottom plate (50 μL cells/well) at a density of 0.5 x 10^6^ cells per well.

### Flow cytometry

Gene editing read-outs were carried out via flow cytometry on 96-well round-bottom plates using a CytoFLEX LX device (Beckman Coulter). In order to allow ample protein turnover and ensure the visualization of transgene expression, cell counting and assessment of knock-in efficiency were performed at 4- and 7-days following electroporation. All staining panels are specified in Supplementary Table 4. Cell concentrations were assessed by acquiring 30 μL of resuspended cells diluted 1:5 in PBS-4′,6-diamidino-2-phenylindole (DAPI) (Thermo Fisher Scientific) without any prior washing steps. For detection of transgene expression, a series of consecutive washing and staining steps were performed. A volume of 40-100 μl of cell suspension was transferred to a 96-well round-bottom plate and rinsed with 100-160 μl of PBS at 400 x g for 5 minutes. Subsequently, the supernatant was removed, and the cell pellets were briefly vortexed. For cells electroporated with HDRTs lacking a Myc tag, an initial surface stain was done using an anti-Fc antibody (αFc) (polyclonal, Jackson Immuno Research) conjugated to an Alexa Fluor 647 (AF647) fluorochrome. To achieve this, 30 μL of diluted αFc were added per staining condition, followed by incubation of cells at 4°C in the dark for 15 minutes. Afterwards, the cells were washed again and stained a second time with a Pacific-Blue-conjugated anti-CD3 antibody (clone UCHT1, Biolegend) and a LIVE/DEAD-UV dye (Thermo Fisher Scientific). As for the cells electroporated with HDRTs containing a Myc tag, CAR detection was achieved with one staining using an anti-Myc antibody (clone 9B11, Cell Signaling Technology) coupled to AF647 and the same anti-CD3 antibody and LIVE/DEAD-UV dye mentioned before.

### *In vitro* AsCas12a cleavage assay with gel electrophoresis read-out

Standard *in vitro* cleavage reactions were conducted using either commercially purchased NEB2 buffer (New England Biolabs) or an in-house-made version composed of 50 mM NaCl, 10 mM Tris-HCl (pH 7.9), 100 μg/μL BSA, with varying concentrations of MgCl_2_ (ranging from 0 to 10 mM). All reactions, containing either buffer, were quenched by the addition of 8-fold molar excess of EDTA (0.5 M, Thermo Fisher Scientific) relative to the highest Mg^2+^ concentration and then loaded on an agarose gel for assessment of ssDNA cleavage. In the assay, 0.5 μL of ssCTS substrates (0.25 μg per condition) were incubated at 37°C for 30 minutes with different reagents depending on the condition tested. AsCas12a (64 μM) and crRNA (100 μM) were diluted 1:10 in nuclease-free water, and volumes corresponding to those from the electroporation were utilized (0.48 μL crRNA and 0.4 μL AsCas12a). All reactions were carried out in a total volume of 5 μL (excluding the EDTA used for reaction quenching), with nuclease-free water added as necessary to achieve this volume.

### Fluorescence-based CRISPR detection assay

Synthetic DNA was detected in a 20 μL CRISPR reaction in a 384-well microplate at 37°C (43). CRISPR detection was performed with final concentrations of 250 nM poly-TTATT-HEX reporter, 90 nM AsCas12a, 45 nM crRNA, 50 mM NaCl, 10 mM Tris-HCl, 100 µg/ml BSA, and either 0, 0.5, 1, 5, 10 mM MgCl_2_. Fluorescence was measured on a multi-mode microplate reader (iD5) with an excitation/ emission wavelength pair of 530/ 570 nm. Fluorescence measurements were read for 90 mins at 5-minute intervals.

### Data analysis, statistics, and presentation

Flow cytometry data were analyzed using FlowJo software version 10 (BD Biosciences). Data from various assays were organized in Excel (Microsoft), and graphs were generated using Prism 9 (GraphPad). The impact of various template formats on gene editing efficacy was assessed through either one-way or two-way repeated measures ANOVA, followed by a Dunn’s correction for multiple comparisons (p < 0.05). Diagrams depicting nucleic acid sequences, receptors, and experimental workflows in the figures were generated using www.biorender.com.

## 4. Results

### Flanking truncated Cas9-target sequences enhance ssDNA-mediated CD19-CAR insertion at the *TRAC* locus

To validate reported effects of tCTS-modified ssDNA HDRTs in primary human T cells, we performed non-viral knock-in experiments with a 2-kb-sized second-generation CD19-CAR (2.8 kb including homology arms (HAs)) for targeted integration into the *TRAC* locus using the previously described SpCas9 CTS design (31). For this, we generated ssDNA from the sense (+) and antisense (-) strand and incorporated tCTS on either the 5’ or 5’ and 3’ ends (Suppl. Fig. 1A). The CTS motifs included a 4-bp buffer sequence, a truncated sgRNA target sequence (with 6-bp mismatches (mm)) and a PAM ‘In’ orientation (facing towards the insert). Prior to electroporation, corresponding oligodeoxynucleotides (ODN) were hybridized to the CTS to create dsDNA ends. Using our CRISPR-Cas9 gene editing protocol without any HDR enhancers (27), we examined the impact of various ssCTS designs on gene insertion rates and the number of transgene-expressing T cells 4 days post-electroporation with 0.5 µg of DNA using flow cytometry (Suppl. Fig. 1B). Consistent with prior findings (31), ssCTS HDRTs performed better than non-modified ssDNA, likely attributed to enhanced CTS-facilitated DNA nuclear delivery (Suppl. Fig. 1C). In contrast to the previous report, we found that modifications both at the 5’ and 3’ end yielded higher HDR frequencies than ssDNA-HDRTs with just a single 5’-tCTS (Suppl. Fig. 1C). Furthermore, compared to conventional dsDNA templates, ssCTS HDRTs with flanking modifications demonstrated comparable HDR efficiencies and knock-in cell counts, with antisense ssCTS yielding higher HDR frequencies and knock-in cell numbers than sense ssCTS. Taken together, the generation of ssDNA from the antisense strand and the inclusion of CTS modifications on both DNA ends resulted in the highest knock-in rates (mean 23% ± SD 10.5) with the SpCas9-nuclease.

### Reduced crRNA mismatches and PAM ‘In’ orientation of Cas12a-target sequence motifs enhance dsCTS knock-in efficacy

We reasoned that the same strategy of using CTS to enhance SpCas9 editing (23) could be adapted to the AsCas12a nuclease. To this end, we first set out to test different CTS configurations using dsDNA HDRTs. We designed CTS motifs specific for Cas12a that included a 16-bp buffer sequence, a PAM, and either an intact or truncated crRNA target sequence (Fig. 1A). Given the previously reported low intrinsic tolerance of AsCas12a for mismatches within regions proximal (1 to 18 bp) to the PAM (37, 40), we tested crRNA target sequences with 0, 2, 4, 6, 8, and 12-bp mismatches. Moreover, we investigated the impact of the orientation of the gRNA-recognition sequence by placing the PAM either ‘In’ (red) or ‘Out’ (blue) of the templates (facing inwards or outwards in relation to the insert). The impact of the CTS motifs on knock-in efficiency was then compared in terms of the relative HDR frequency (measured by flow cytometry) and the number of transgene-expressing cells relative to unmodified dsDNA (Fig. 1B). To exclude constructor locus-specific bias, we conducted a screening of the different CTS motifs in HDRTs for three different knock-in strategies, designed to introduce CAR transgenes of different sizes at three distinct loci of the T-cell receptor complex. These included a 0.8kb-sized HLA-A2-specific TRuC at the *CD3*ε (smallest transgene), a 1kb-sized truncated CD19-CAR at the *CD3*ζ-gene or a complete 2kb-sized CD19-CAR the *TRAC-* locus (largest transgene) (Fig. 1C).

**Figure 1:**
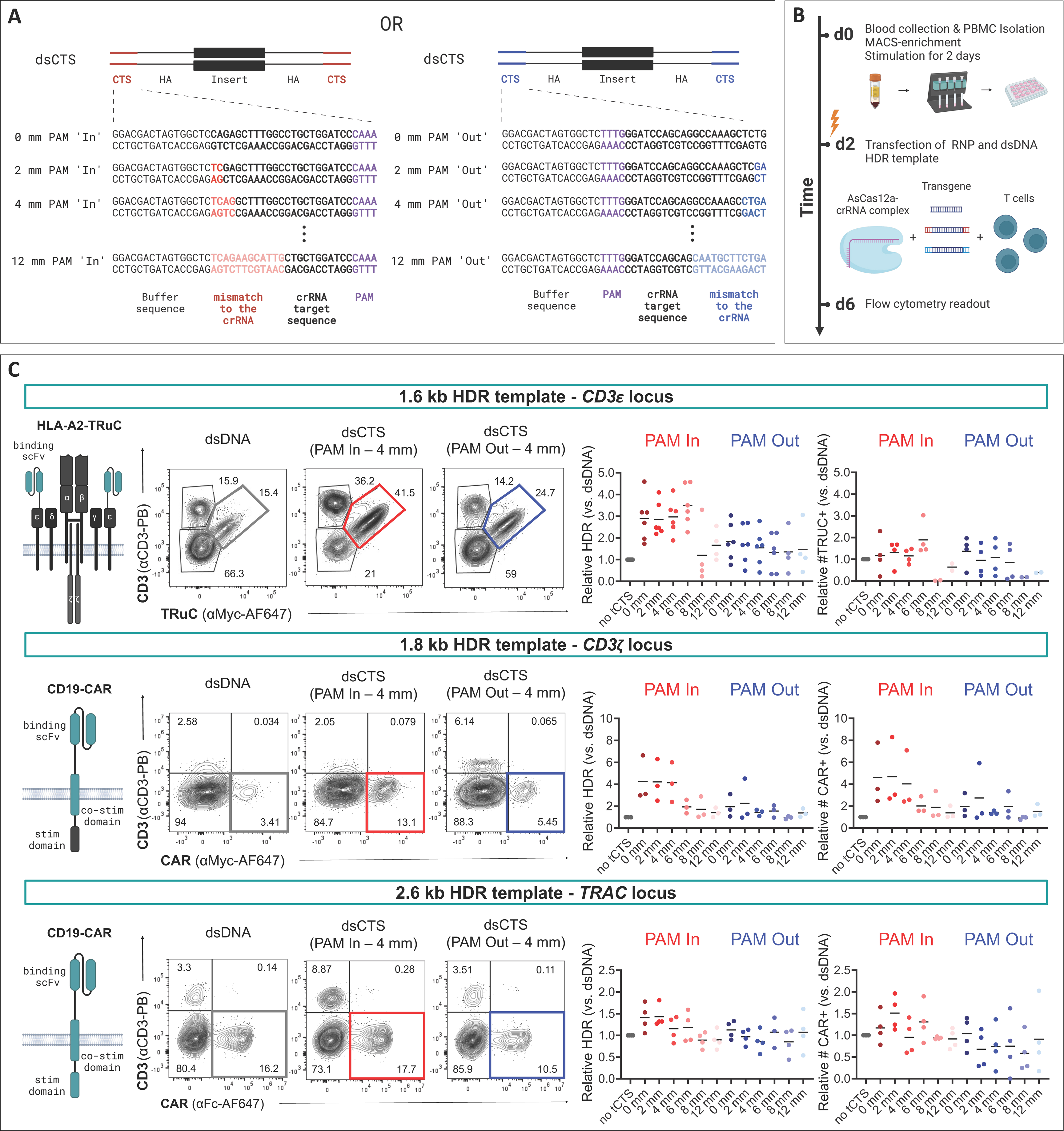
Cas12a-target sequence-motifs with fewer crRNA-mismatches and PAM ‘In’ orientation increase knock-in efficiency of dsDNA CAR or TRuC constructs independent of the edited locus. Virus-free insertion of an HLA-A2-TRuC and two CD19-CAR transgenes into the human *TRAC*, *CD3*ζ, or *CD3*ε loci. (A) Designs of dsCTS donor templates are shown. The inserts are flanked by HAs with additional AsCas12a CTS. These include a 16-bp buffer region, a PAM, and either an intact (CTS with 0-bp mm) or truncated crRNA sequence (tCTS with 2, 4, 6, 8 or 12-bp mm). Based on the PAM orientation, templates are referred to as PAM ‘In’ (3’ of the crRNA sequence, red) or PAM ‘Out’ (5’ of the crRNA sequence, blue). (B) Experimental setup to evaluate co-electroporation of RNPs and dsDNA donor templates with or without CTS motifs. (C) Left side, schematics of transgene-encoded receptors and representative flow cytometry plots depicting editing outcomes using non-modified dsDNA HDRTs, or with 4-bp mm PAM ‘In’ or PAM ‘Out’. Right side, Summary of flow cytometric analysis 4 days after electroporation (n = 6 for *CD3*ε-, n = 3 for *CD3*ζ- and n = 4 for *TRAC*-knock-in into healthy donors). Black lines indicate mean values. HDR efficiencies and number of transgene-expressing cells are shown relative to the dsDNA condition in grey. For each knock-in condition, 0.5 μg template was used.

With dsDNA HDRTs, CTS modifications with PAM ‘In’ and lower number of mm increased the relative knock-in rates over unmodified dsDNA HDRTs. The relative increases in HDR frequencies were more evident for smaller than for the largest HDRT: For the *CD3*ε-directed HDRT containing the smallest insert (0.8kb-HLA-A2-TRuC), the inclusion of CTS with PAM ‘In’ (and up to 6 mms) resulted in up to a 4.5-fold increase in HDR efficiency of dsCTS relative to dsDNA (Fig. 1C, top panel). Similarly, for the *CD3*ζ-directed 1-kb-CD19-CAR HDRT (1.8 kb including HAs), the addition of CTS PAM ‘In’ also enhanced HDR efficacy in some conditions, such as CTS PAM ‘In’ 0, 2, and 4-bp mm (Fig. 1C, middle panel). For instance, templates with a 4-bp-mm CTS PAM ‘In’ demonstrated on average a 4-fold increase (± SD 1.48) in HDR frequencies compared to dsDNA. In the case of the *TRAC*-directed 1.8-kb-CD19-CAR (2.6 kb including HAs), increased knock-in rates were less pronounced and were only observed with CTS PAM ‘In’ templates with the least number of mm (Fig. 1C, bottom panel). For example, dsCTS with a 2-bp mm and a PAM ‘In’ orientation showed on average an increase in HDR of 1.4-fold (± SD 0.2). The number of transgene-positive T cells was only increased in conditions with the *CD3*ζ-HDRT with CTS PAM ‘In’ and few mismatches. In all other conditions and HDRTs, the relative increases in HDR frequencies did not result in higher edited T cell yields (Fig. 1C). Overall, relative improvements of CRISPR-Cas12a-mediated knock-in rates were most evident when utilizing templates with fewer crRNA mm ranging 0 to 4 and a PAM ‘In’ orientation in the CTS region. These designs were further evaluated in the ssCTS format.

### Flanking double-stranded CTS improve HDR efficiency of ssDNA without inducing ssDNase activity of AsCas12a under physiological magnesium concentrations

Inclusion of dsDNA CTS end modification in ssDNA HDRTs could trigger unspecific DNase activity after binding of the AsCas12a-crRNA complex to its target (the CTS) *in vitro* (prior to electroporation into the T cells) or *in cellulo* (after electroporation). To evaluate the propensity of AsCas12a Ultra to degrade ssDNA HDRTs with CTS *in vitro*, we generated ssDNA from the antisense strand and incorporated CTS on both ends (Fig. 2A). The CTS motifs included a 4-bp buffer sequence, a complete or 4-bp mm crRNA target sequence and a PAM ‘In’ ori-entation. To assess whether AsCas12a indiscriminately degrades ssCTS templates, the HDRTs were co-incubated with the crRNA-nuclease complex employed in electroporation experiments. This was performed at 37°C for 30 minutes with or without the NEB2 buffer containing a high concentration of Mg^2+^, which served as a positive control since it activates Cas12 to indiscriminately degrade ssDNA even without the presence of a crRNA (41) (Fig. 2B, left side). Non-specific nuclease activity was detected solely in the presence of high Mg^2+^, independent of the crRNA (Fig. 2B, right side). Moreover, the inclusion of either complete or truncated CTS did not affect the nonspecific cleavage activity of AsCas12a. Given the high Mg^2+^ level in the NEB2 buffer (10 mM), we hypothesized that no degradation of ssCTS would occur under lower Mg^2+^ conditions that mimic physiological intracellular concentrations (0.2 - 1 mM). To test this, we co-incubated the HDRTs with the crRNA-nuclease complex in an in-house prepared NEB2-like buffer containing a range of Mg^2+^ concentrations. Comparable to the buffer-free condition, there was no evident nonspecific degradation of either intact or truncated ssCTS in the presence of the buffer containing low Mg^2+^ concentrations (Fig. 2C). To ensure the feasibility of proceeding with this nuclease in gene editing experiments, we decided to perform a more sensitive digestion assay (43), that detects *trans*-cleavage of ssDNA reporters as an indicator of AsCas12a collateral ssDNase activity (Fig. 2D). To mimic an intracellular setting, we conducted this assay in buffers with varying Mg^2+^ concentrations. As expected from the previous digestion assays, *trans*-cleavage activity was detected only when high Mg^2+^ concentrations were used. When comparing the two ssCTS templates, there was no difference in the measured *trans*-cleavage under low Mg^2+^ levels, as no digestion occurred.

**Figure 2:**
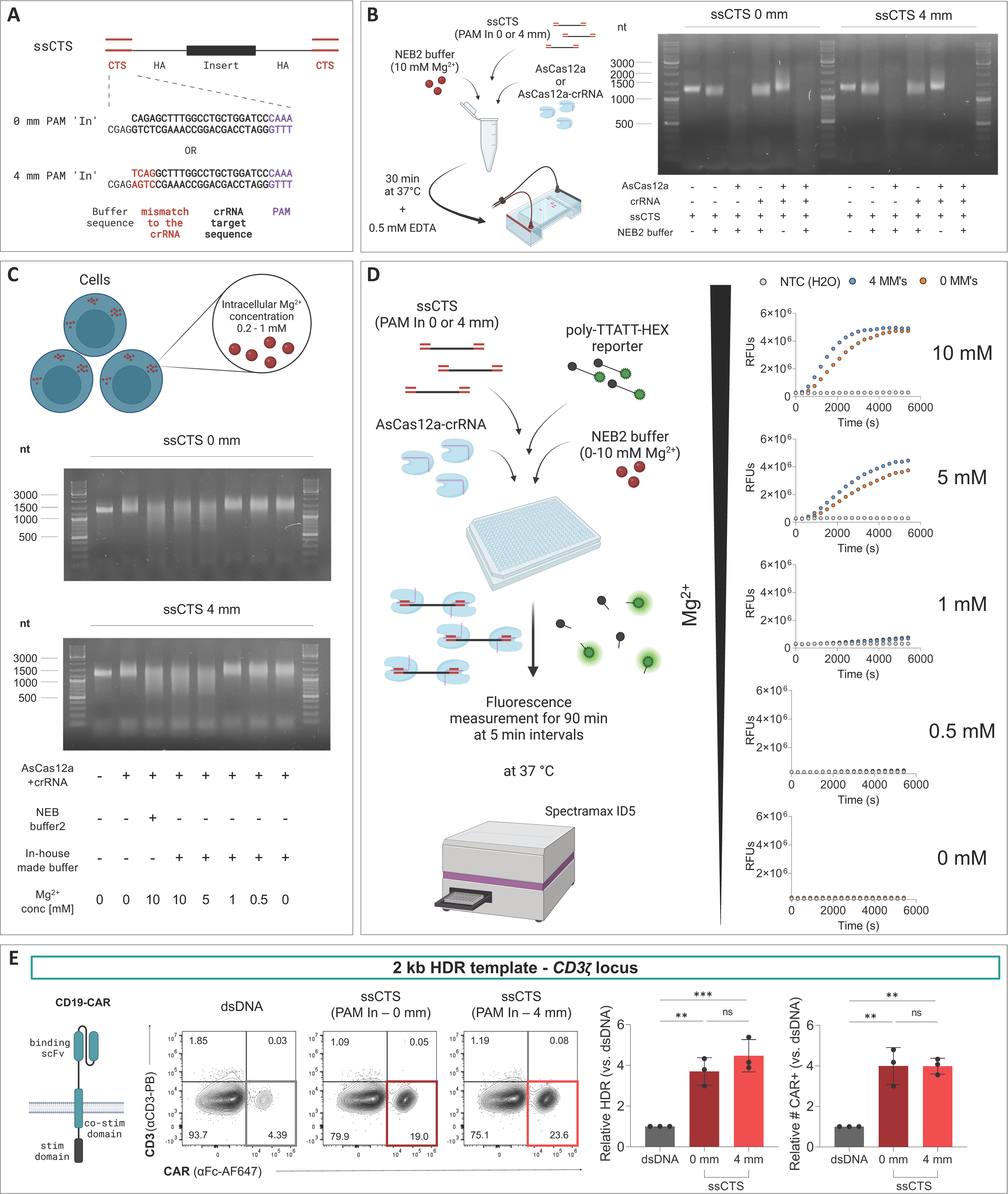
Hybrid dsDNA modifications with Cas12a-target sequence improve knock-in efficacy of linear ssDNA without triggering ssDNase activity of AsCas12a under physiological Mg^2+^concentrations. (A) Designs of ssCTS donor templates are shown. The insert is flanked by HAs and hybridized CTS motifs. The CTS includes a 4-bp buffer region on the antisense strand, a PAM and either an intact (CTS with 0-bp mismatches) or truncated crRNA sequence (tCTS with 4-bp mm). (B) *In vitro* Cas12a cleavage assay depicting digestion of both intact and truncated ssCTS templates in 10 mM Mg^2+^-containing NEB2 buffer after 30 minutes incubation at 37°C followed by reaction quenching with 0.5 mM EDTA. (C) *In vitro* Cas12a cleavage assay depicting digestion of both intact and truncated ssCTS templates in either 10 mM Mg^2+^-containing NEB2 buffer or an in-house prepared buffer with various Mg^2+^concentrations (absence, physiological intracellular conditions 0.2-1 mM, or excess 5-10 mM) after 30 minutes incubation at 37°C and reaction quenching with 0.5 mM EDTA. (D) Fluorescence-based CRISPR detection assay utilizing a poly-TTATT-HEX reporter was employed for detection of Cas12a-cleavage activity across varying Mg^2+^-concentrations (0, 0.5, 1, 5, and 10 mM Mg^2+^). Fluorescence measurements were acquired for 90 minutes at 5minute intervals using a multi-mode microplate reader (Spectramax iD5) with an excitation/emission wavelength pair of 530/570 nm. (E) Left side, schematic of CD19-CAR receptor and representative flow cytometry plots depicting editing outcomes using dsDNA and intact or 4-bp mm ssCTS PAM ‘In’. Right side, Summary of flow cytometric analyses 4 days after electroporation (n = 3 for *CD3*ζ-knock-in into healthy donors). Thick lines indicate mean values; error bars indicate standard deviation. Black dots represent individual data points. HDR efficiency and numbers of CAR-expressing cells are shown relative to the dsDNA condition in grey. For each knock-in condition, 0.5 μg template was used. Statistical analysis was performed using ordinary one-way ANOVA with subsequent Dunn’s correction (for multiple testing) comparing values for each HDRT format with dsDNA as reference. Asterisks represent different p values calculated in the respective statistical test (not significant [ns]: p > 0.5; ∗p < 0.05; ∗∗p < 0.01; ∗∗∗p < 0.001).

After excluding a ssDNase activity of AsCas12a in Mg^2+^-low environments, we proceeded to test the intact and 4-bp truncated ssCTS HDRTs in electroporation experiments using the same workflow as previously outlined (Fig. 1B). The use of ssCTS templates led to a relative increase in knock-in rates by at least 3-fold (up to 5.5-fold). In contrast to previous experiments with dsCTS HDRTs (Fig. 1), ssCTS also increased the absolute number of CAR-expressing cells relative to non-modified dsDNA (Fig. 2E). When comparing intact versus truncated ssCTS, slightly higher HDR rates were observed when a 4-bp mm to the crRNA was included in the CTS motifs, but these differences were not statistically significant. These results suggest that both intact and truncated ssCTS can be utilized for efficient AsCas12a Ultra-mediated HDR, achieving significantly higher HDR rates and number of CAR-expressing cells compared to dsDNA templates.

### Incorporating a buffer region and introducing crRNA-mismatches into the CTS region enhance gene editing outcome with AsCas12a

After confirming the efficacy of ssCTS as a template for CRISPR-Cas12a gene editing, we aimed to further optimize the CTS motifs for ssDNA. To this end, we evaluated the impact of different buffer sequences adjacent to the CTS (Suppl. Fig. 2A). We created ssCTS either with or without a 4-bp buffer sequence placed on the template strand alone (OS) or on both the template strand and the annealed ODN (TS) (Suppl. Fig. 2A). The inserted template for gene editing at the *CD3*ζ locus was the same 1.2-kb-sized CD19-CAR (2 kb including HAs) as used before (Fig. 2E). When comparing ssCTS without a buffer sequence, there was no statistically significant difference to the non-modified ssDNA conditions (Suppl. Fig. 2B). In contrast, higher HDR frequencies were observed with the addition of an OS or TS-buffer, especially when using truncated templates. For instance, the transgene was detected in an average of 22% of T cells (± SD 8.25) when using OS ssCTS 2 mm, and in 21% of T cells (± SD 8.76) with OS ssCTS 4 mm. Given the similar performance of these formats, we proceeded with OS 4 mm in the subsequent experiments.

### ssCTS donors outperform dsDNA templates at high concentrations independent of the transgene or insertion site

Following the selection of the most optimal CTS modification for CRISPR-Cas12a gene editing, we aimed to compare our ssCTS template with dsDNA, dsCTS, and ssDNA. The Cas12a-binding motifs of both ssCTS and dsCTS included a buffer sequence, a 4-bp mm to the crRNA and a PAM ‘In’ orientation (Fig. 3A). To eliminate any bias specific to the construct or locus, we compared different concentrations of the various HDRT formats by insertion of an HLA-A2-TRuC into the *CD3*ε locus and CD19-CARs into *CD3*ζ or *TRAC* locus. In the case of the HLA-A2-TruC (1.6 kb including HAs), the incorporation of CTS motifs increased HDR of dsCTS and ssCTS compared to dsDNA and ssDNA, respectively, across various concentrations (Fig. 3B, top panel). However, with dsCTS, toxic doses were encountered starting at 50 nM, resulting in a progressive decline in knock-in efficiency and total cell yield. In contrast, ssCTS templates did not affect cell yield, even at the highest tested concentration (100 nM). Additionally, HDR frequencies increased with higher template concentrations without reaching a plateau. Notably, transgene expression was detected in up to 90% of T cells when using ssCTS at the highest HDRT concentration. When repeated independently in another laboratory using the same batch of ssCTS HDRT or a newly generated batch, HDR efficiencies were comparably high, reaching up to 92% (Suppl. Fig. 3). Similar trends were observed for *CD3*ζ-directed HDR insertion of a CD19-CAR, with ssCTS templates leading to transgene detection in up to 44% of T cells at the highest concentration (Fig. 3B, middle panel). Lastly, non-viral gene editing was conducted using the larger *TRAC*-CD19-CAR knock-in construct (2.6 kb including HAs) (Fig. 3B, lower panel). Both dsDNA and dsCTS templates exhibited high frequencies of HDR at concentrations of 25 nM to 50 nM but also showed a decrease in efficiency concomitant with a loss in cell numbers. In contrast, ssCTS showed a gradual increase in HDR, achieving transgene insertion frequencies ranging from 5% at 12.5 nM to 35% at 100 nM without a drop in cell numbers.

**Figure 3:**
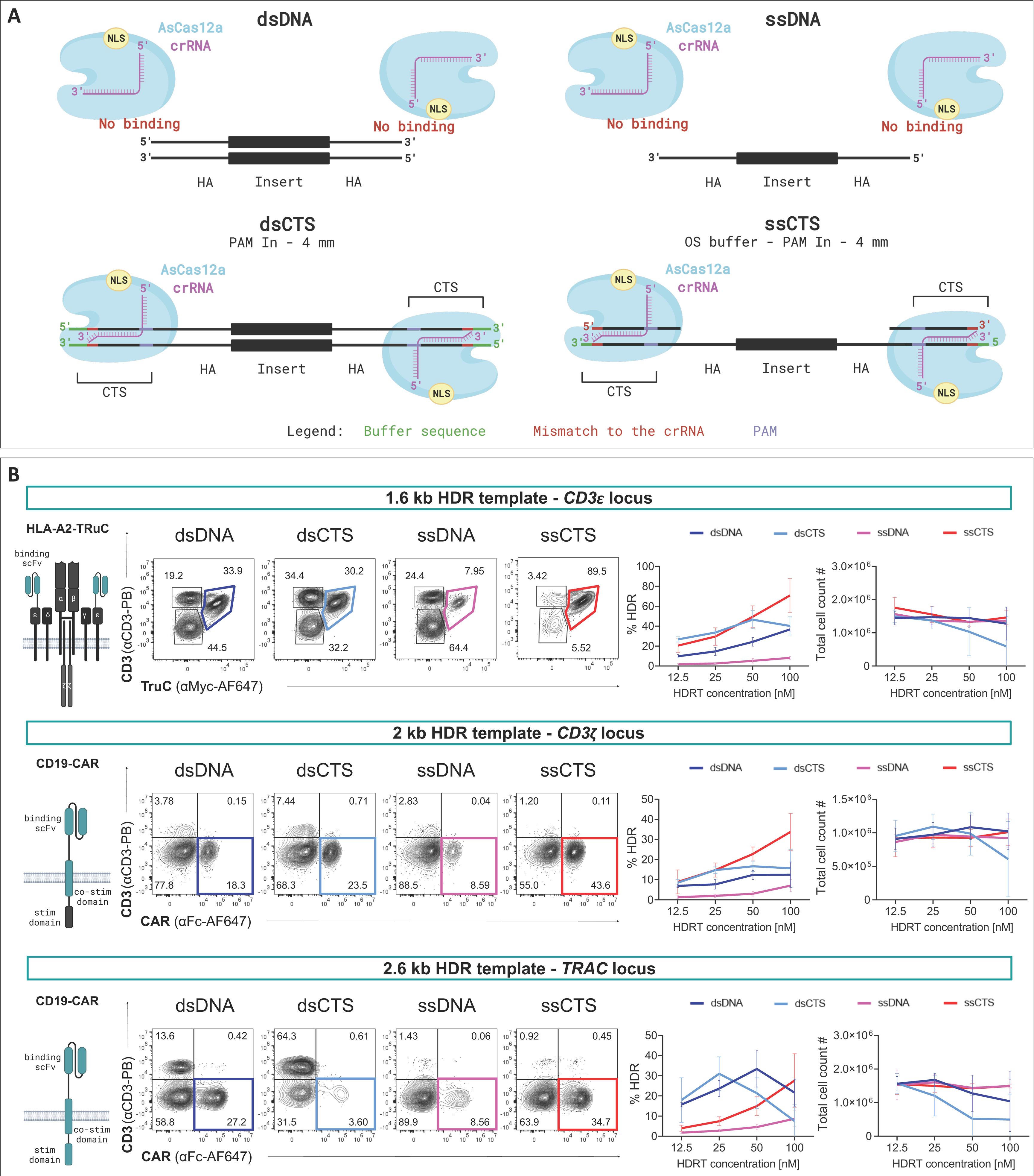
Truncated CTS-modified ssDNA containing a buffer sequence and PAM ‘In’ orientation outperform dsDNA templates at high concentrations. Virus-free insertion of an HLA-A2-TRuC or two CD19-CAR transgenes into the human *TRAC*, *CD3*ζ or *CD3*ε loci. (A) Schematics of the interactions of the NLS-containing Cas12a-crRNA complex with the CTS motifs of dsDNA and ssDNA templates. Both CTS-containing construct-formats contain CTS truncated by a 4-bp mm, have a PAM ‘In’ orientation and a buffer region. (B) Left side, schematics of transgene-encoded receptors and representative flow cytometry plots depicting editing outcomes using dsDNA, dsCTS, ssDNA, and ssCTS HDRTs. Right side, Summary of flow cytometric analyses 4 days after electroporation (n = 3 for *CD3*ε-, n = 4 for *CD3*ζ- and n = 3 for *TRAC*-knock-in into healthy donors, each from 2 independent experiments). Dark and light blue indicate the use of dsDNA and dsCTS templates, whilst pink and red depict ssDNA and ssCTS, respectively. Thick lines indicate mean values; error bars indicate standard deviation. The percentage of HDR efficiency and total cell numbers are shown in a HDRT concentration-dependent manner.

In summary, irrespective of the transgene or target locus, CTS motifs were essential additions to ssDNA templates to facilitate highly efficient CRISPR-Cas12a gene knock-in. With these end modifications, high concentrations of ssCTS consistently resulted in higher knockin rates and improved cell yields compared to dsDNA and dsCTS.

## 5. Discussion

The field of gene therapy has been rapidly progressing, marked by numerous ongoing clinical trials and extensive preclinical studies exploring innovative products (44). CRISPR-Cas technology is gaining prominence due to its potential to tackle existing safety concerns and simplify the complex and costly manufacturing processes of gene therapy products, such as CAR-T cells (3). However, further optimization is necessary to maximize the yield and improve the quality of gene-modified cells. One defining parameter determining the efficiency of gene editing is the donor template format. Despite the current scarcity of systematic studies comparing the design elements of synthetic templates, it has been established that ssDNA constructs are notably less harmful to cells (15). Here, we demonstrated that the manufacture of genetically engineered T cells using CRISPR-Cas12a could be significantly improved by employing ssDNA templates with double-stranded CTS end modifications.

Our results suggest that incorporating CTS into linear ssDNA donor templates is a powerful tool to optimize gene transfer with different Cas species and in a non-viral fashion. We have independently replicated findings by Shy et al. (31) reporting that ssCTS templates significantly improve HDR efficiency with SpCas9-mediated gene editing in comparison to dsDNA or unmodified ssDNA. To translate the ssCTS approach to another Cas editing platform, we reasoned that optimization of CTS would be necessary to account for the differences in target interrogation between orthogonal Cas enzymes. This involved examining dsDNA containing both intact and truncated CTS sequences. Similar to previously reported CTS modifications for SpCas9 (23, 31), PAM ‘in’ orientation and low number of mismatches up to 4 bp at the 5’ end of the crRNA target sequence resulted in enhanced HDR efficiencies with the AsCas12a Ultra enzyme. For SpCas9, truncated CTS was superior to CTS without mm. With AsCas12a, constructs with perfect CTS lacking any mismatches but containing a PAM 3’ of the crRNA (PAM ‘In’) also outperformed non-modified donor templates. This may be attributed to the characteristic of Cas12a to cut distally from the PAM, thereby restricting the cleavage to the 5’ end of the template. Moreover, excessive mm to the crRNA-target sequence led to a decrease in editing efficiency, likely due to the reported low intrinsic tolerance of AsCas12a for mismatches within regions proximal (1 to 18 bp) to the PAM (37, 45). These results increase the confidence that incorporation of CTS can be adapted to other Cas species, beyond SpCas9 and AsCas12a Ultra.

To utilize the AsCas12a nuclease for gene editing with ssCTS templates, we investigated its reported non-specific ssDNase activity as previously described (41, 42). Collateral Cas12a-mediated damage to ssDNA HDRTs after CTS-binding *in vitro* or induction of the DSB at the on-target locus would be detrimental to efficient gene editing. Our original hypothesis was that templates with double-stranded truncated CTS would be less susceptible to digestion compared to intact ones. However, cleavage of ssCTS templates occurred only under conditions with non-physiologically high concentrations of Mg^2+^, confirming previous findings (41). Considering that magnesium is the fourth most abundant positively charged ion in the body and the second most abundant within cells (46), this raised questions about how intracellular magnesium levels might influence the nonspecific cleavage of ssCTS by AsCas12a. As the concentration of free magnesium inside cells varies between 0.5 and 1 mM (47), we aimed to investigate ssDNase activity under physiological Mg^2+^ concentrations. By measuring *cis*- or *trans*-cleavage under varying Mg^2+^ concentrations and testing the nuclease with our templates from electroporation experiments, our findings indicated that AsCas12a does not induce excessive ssDNA degradation. Consequently, no discernible difference was observed between the intact and truncated CTS constructs. Our results suggest that there is no overt ssDNase activity of AsCas12a Ultra in human T cells. Others have previously reported that ssDNase activity of Cas12a systems did not contribute to bacterial host defense against bacteriophages (48), suggesting that the ssDNase phenomenon could be primarily restricted to *in vitro* settings. Moreover, one study demonstrated that *in vitro* collateral ssDNase activity is less pronounced in AsCas12a than in other Cas12a enzymes such as LbCas12a(49).

The ssCTS templates enabled efficient gene transfer of various clinically relevant, intermediate-sized transgenes, but with notable template-size dependent variabilities. The ssCTS templates outperformed dsCTS HDRTs in terms of both cell viability and HDR efficacy at the highest concentration tested. In general, notable differences were observed in the transgene insertion efficiency between larger and smaller constructs. For instance, using a 0.8 kb-sized insert into the *CD3*ε gene led to the detection of the transgene in up to 90% of the cells, whereas only 35% of the cells expressed a 1.8 kb-sized insert at the *TRAC* locus. While these differences might also be locus-dependent, the observed association between size and toxicity is consistent with previous findings with the SpCas9-ssCTS platform (31), suggesting that larger HDRTs tend to induce greater toxicity in cells. This toxic effect was considerably more pronounced in dsDNA templates compared to ssCTS. The exact reason for the reduced toxicity of ssCTS relative to dsDNA in cellular systems is not yet fully elucidated, but it may be attributed to differential recognition by DNA-sensing pathways (50, 51) or to increased physical stress by the larger dsDNA-RNP aggregates. In our experiments, we did not reach a concentration at which ssCTS templates induced dose-dependent toxicity. As a next step, investigating higher concentrations of larger-sized ssCTS would be valuable to determine the highest potential knock-in rate for larger constructs. However, the production of highly concentrated and pure linear ssDNA remains a limitation to execute these suggested experiments. In our hands, linear ssDNA HDRTs production by single-strand exonuclease digestion of dsDNA (see methods) becomes very inefficient for constructs larger than 3kb. Other methods include biotin-streptavidin bead selection of a labelled DNA strand (31, 52) or asymmetric PCR (53), but they suffer from reduced purity. Alternatively, others have demonstrated that circular ssDNA produced from phagemids is suitable for large-scale production of large HDRTs, up to 13 kb in size (54). Future studies may investigate whether knock-in efficacy of circular ssDNA HDRTs can be increased with CTS.

The experiments with ssCTS with SpCas9 (31) and AsCas12a (this study) demonstrate that insufficient nuclear delivery after electroporation of non-viral ssDNA templates likely impedes efficient editing. Alternative strategies to increase nuclear concentration of ssDNA HDRTs involve other means of coupling the DNA donors with the NLS-tagged gene editor directly, creating a tripartite complex of NLS-SpCas9 with sgRNA and circular ssDNA templates (55). Future studies may elucidate other means to deliver ssDNA templates for efficient, non-toxic editing and explore different transfection modalities, such as lipid nanoparticles or chemical transfectants, with and without peptide-mediated Cas delivery (56, 57).

Altogether, by incorporating CTS to overcome the nuclear transport barrier and by leveraging the decreased cellular toxicity of ssDNA, ssCTS constructs provide a compelling alternative to dsDNA-mediated HDR in non-viral CRISPR-Cas12a gene editing. For smaller transgenes, such as CD3ε-TRuC used in our study, ssCTS templates can reach integration rates as high as 90%. These efficiencies mirror integration frequencies previously only achieved with recombinant adeno-associated virus for template delivery (58). Moreover, ssCTS templates offer a promising solution to the hurdle of obtaining sufficient numbers of transgene-positive cells without the need for additional purification steps (18), pharmacological enhancers (27, 31), or knock-in into essential genes (59) since higher donor concentrations can be used without causing relevant toxicity. The AsCas12a-ssCTS platform contributes to improving the virus-free manufacturing process for adoptive T-cell-based therapies for future clinical applications.

## 7. Supplementary Figure Legends

**Supplementary Figure 1:**
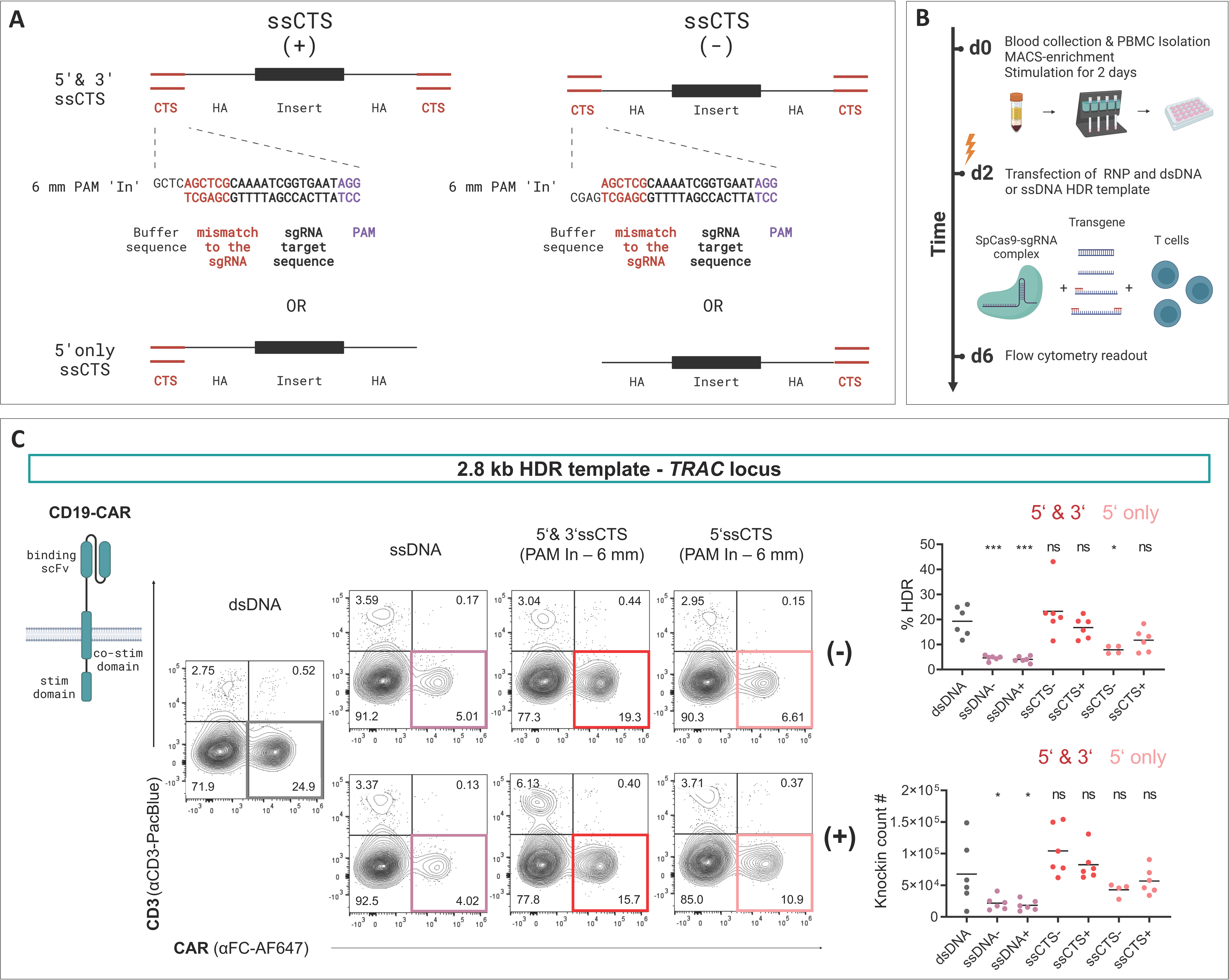
Truncated Cas9-target sequence-motifs with PAM ‘In’ orientation on both ssDNA ends increase knock-in efficiency of CD19-CAR construct at the TRAC locus. (A) Designs of ssCTS donor templates are shown. The inserts are flanked by HAs with additional SpCas9 CTS on one (5’ only) or both ends (5’ & 3’) of the constructs. The CTS include a 4-bp buffer region, a PAM ‘In’ orientation, and a truncated crRNA sequence (tCTS with 6-bp mm). Based on the remaining strand after enzymatic digestion, ssCTS templates are referred to as “(+)” or “(-)“. (B) Experimental setup to evaluate coelectroporation of RNPs and the different ssCTS constructs versus dsDNA. (C) Left side, schematic of the expressed CD19-CAR and representative flow cytometry plots depicting editing outcomes using dsDNA, ssDNA without CTS or with 6-bp tCTS on the 5’ and 3’ ends or 5’ end only. Right side, summary of flow cytometric analysis 4 days after electroporation (n = 7 for TRAC-knock-in into healthy donors). Black lines indicate mean values. Percentage of HDR efficiency and numbers of CAR-expressing cells are shown. For each knock-in condition, 0.5 μg template was used. Statistical analysis was performed using ordinary one-way ANOVA with subsequent Dunn’s correction (for multiple testing) comparing values for each HDRT format with dsDNA as reference. Asterisks represent different p values calculated in the respective statistical test (not significant [ns]: p > 0.5; ∗p < 0.05; ∗∗p < 0.01; ∗∗∗p < 0.001).

**Supplementary Figure 2:**
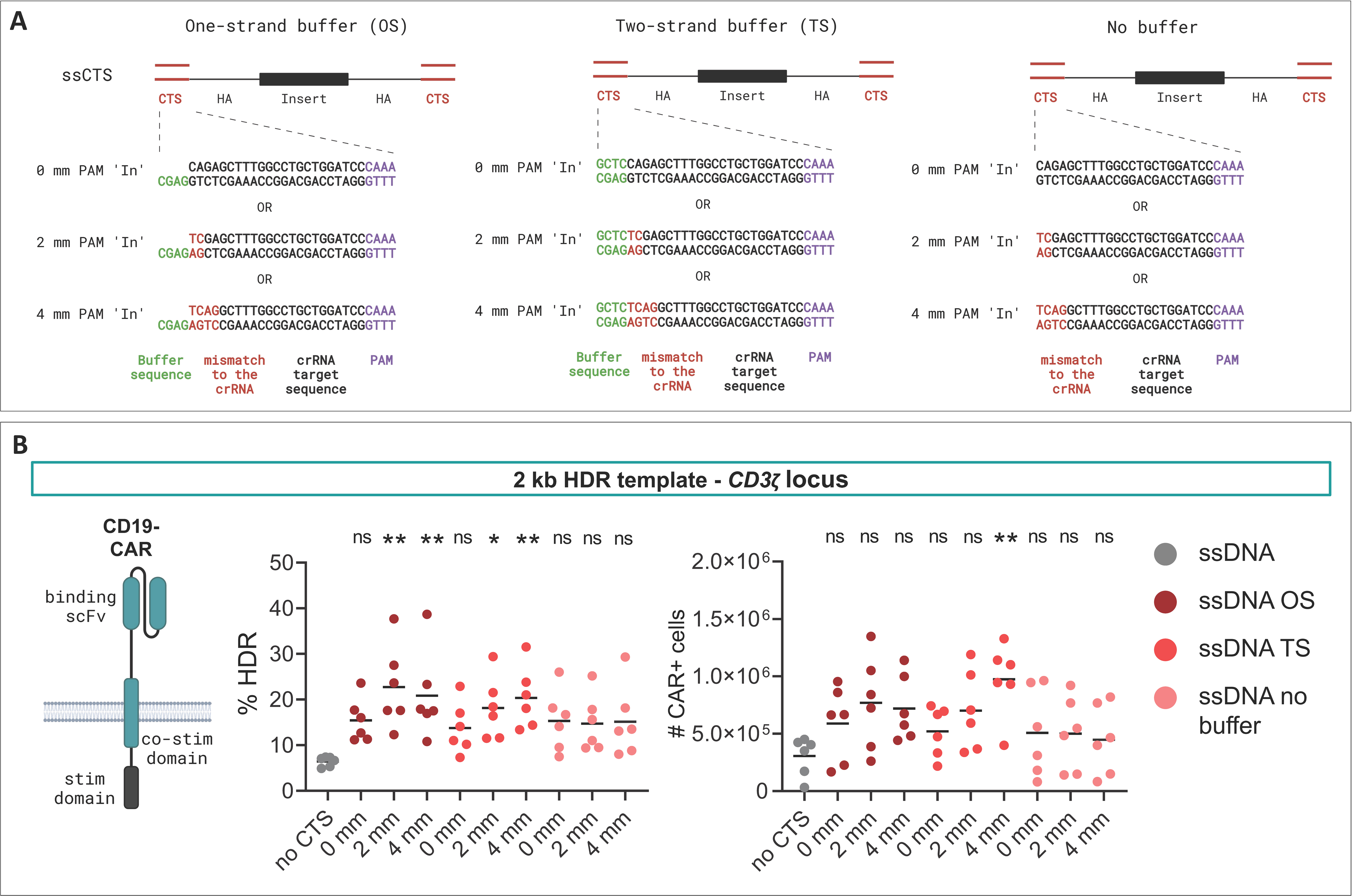
Addition of buffer regions and mismatches to the crRNA in the CTS motifs improve CRISPR-Cas12a gene editing outcome with ssDNA donors. Virus-free insertion of a CD19-CAR transgene into the human CD3ζ locus. (A) Design of ssCTS donor templates is depicted. The insert is flanked by HAs with additional AsCas12a CTS. The CTS comprise a an intact (0-bp mm) or truncated CTS (2- or 4-bp mm), along with a PAM and either no buffer region or a 4-bp buffer region, located on the antisense strand (OS) or on both strands (TS). (B) Left side, schematic of the CD19-CAR receptor. Right side, summary of flow cytometric analyses 4 days after electroporation (n = 6 for CD3ζ-knock-in into healthy donors from 3 independent experiments). Black lines indicate mean values. HDR efficiency and numbers of CAR-expressing cells are shown. For each knock-in condition, 1 μg template was used. Statistical analysis was performed using ordinary one-way ANOVA with subsequent Dunn’s correction (for multiple testing) comparing values for each HDRT format with ssDNA ‘no CTS’ as a reference. Asterisks represent different p values calculated in the respective statistical test (not significant [ns]: p > 0.5; ∗p < 0.05; ∗∗p < 0.01; ∗∗∗p < 0.001).

**Supplementary Figure 3:**
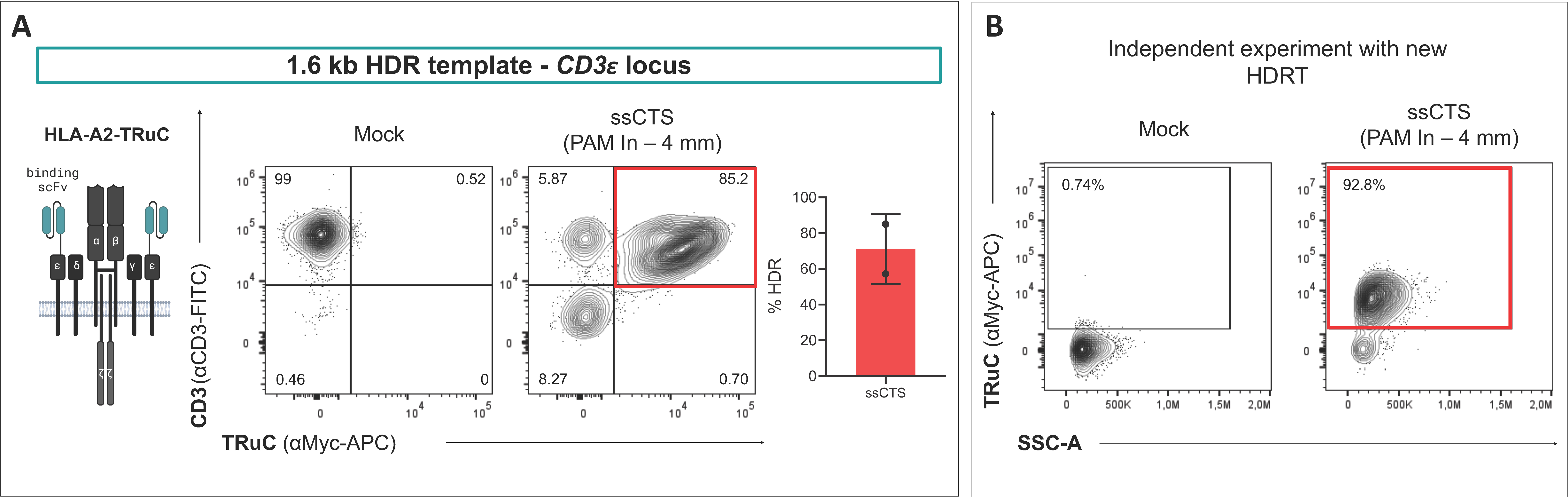
Independent CRISPR-Cas12a knock-in experiments using the HLA-A2-TruC construct confirm the efficacy of the ssCTS templates. Virus-free insertion of an HLA-A2-TRuC transgene into the human *CD3*ε locus. (A) Left side, schematic of the expressed receptor and representative flow cytometry plots depicting editing outcomes using the ssCTS HDRT with 4-bp mm from figure 3B. Right side, summary of flow cytometric analysis depicting HDR efficiency 4 days after electroporation (n = 2 for CD3ε -knock-in into healthy donors from one experiment). (B) Independent gene editing experiment using a newly generated ssCTS template batch encoding the same transgene as in A. Flow cytometry plots depicting editing outcomes 4 days after electroporation (n = 1 for *CD3*ε-knock-in into a healthy donor).

## Supporting information

Supplementary Data File (Supplementary Figures and Table Description)

## 8. Acknowledgements

The authors would like to thank Tatiana Zittel (Charité) for her technical assistance.

AMN, JK and DLW would like to thank their direct support by the Jonas Center for Cellular Therapy of the University of Chicago, USA. This project has received funding from the European Union under Grant Agreement Nr. 101057438 (geneTIGA: genetiga-horizon.eu) to RB and DLW. Views and opinions expressed are however those of the author(s) only and do not necessarily reflect those of the European Union or the European Health and Digital Executive Agency (HADEA). Neither the European Union nor the granting authority can be held responsible for them. JK and DLW were supported by the SPARK-BIH program by the Berlin Institute of Health, Germany. M.M.K. was supported by the Emmy Noether Programme (grant no. KA5060/1-1) of the German Research Foundation and is a participant in the BIH Charité Clinician Scientist Program funded by the Charité – Universitätsmedizin Berlin and the Berlin Institute of Health at Charité (BIH).

## 9. Author contributions

AMN planned, and performed experiments, analyzed results, interpreted the data, and wrote the manuscript. WD, VG designed the first CTS-templates for AsCas12a editing, planned and performed experiments, analyzed results, interpreted the data, and edited the manuscript. JK performed experiments, advised on the figures, interpreted data and edited the manuscript. MS performed experiments and analyzed results. RG and MK planned, performed, analyzed, and interpreted data from *trans*-cleavage assay. NSM and RB planned, performed, and interpreted replication studies with ssCTS. MK and RB provided reagents. DLW designed and led the study, planned experiments, interpreted data, and drafted figures and edited the manuscript. All authors reviewed, commented, and approved the manuscript in its final form.

## 11. Data Availability Statement

All construct sequences can be found in Supplementary Table 1. The CTS designs and corresponding DNA oligo sequences can be found in Supplementary Table 2. The plasmids encoding the original *CD3*ζ*-*HDRT and the *TRAC-*HDRT are available via Addgene (CD3ζ-truncCARgsg: Addgene ID 215759, TRAC-Cas12a: 215769). All other data can be requested from the corresponding author upon request.

