## Supplementary Data File (Supplementary Figures and Table Description) for "Single-stranded HDR templates with truncated Cas12a binding sequences improve knock-in efficiencies in primary human T cells"

**Supplementary Figures	Page**

2**Supplementary Figure 3: Independent CRISPR-Cas12a knock-in experiments using the HLA-A2-TruC construct confirm the efficacy of the ssCTS templates.**

**Supplementary Figure 1: Truncated Cas9-target sequence-motifs with PAM ‘In’ orientation on both ssDNA ends increase knock-in efficiency of CD19-CAR construct at the *TRAC* locus.**

3 **Supplementary Figure 3: Independent CRISPR-Cas12a knock-in experiments using the HLA-A2-TruC construct confirm the efficacy of the ssCTS templates.**

**Supplementary Figure 2: Addition of buffer regions and mismatches to the crRNA in the CTS motifs improve CRISPR-Cas12a gene editing outcome with ssDNA donors.**

4 **Supplementary Figure 3: Independent CRISPR-Cas12a knock-in experiments using the HLA-A2-TruC construct confirm the efficacy of the ssCTS templates.**

**Supplementary Figure 3: Independent CRISPR-Cas12a knock-in experiments using the HLA-A2-TruC construct confirm the efficacy of the ssCTS templates.**

**Supplementary Tables (see Excel File)**

**Supplementary Table 1: DNA sequences of HDR constructs**

**Supplementary Table 2: Annotated primer sequences for different Cas-target sequences**

**Supplementary Table 3: Guide RNA target sequences**

**Supplementary Table 4: Flow cytometry staining information including antibody details**

**Supplementary Figure 1: Truncated Cas9-target sequence-motifs with PAM ‘In’ orientation on both ssDNA ends increase knock-in efficiency of CD19-CAR construct at the TRAC locus.** (A) Designs of ssCTS donor templates are shown. The inserts are flanked by HAs with additional SpCas9 CTS on one (5’ only) or both ends (5’ & 3’) of the constructs. The CTS include a 4-bp buffer region, a PAM ‘In’ orientation, and a truncated crRNA sequence (tCTS with 6-bp mm). Based on the remaining strand after enzymatic digestion, ssCTS templates are referred to as “(+)” or “(-)“. (B) Experimental setup to evaluate co-electroporation of RNPs and the different ssCTS constructs versus dsDNA. (C) Left side, schematic of the expressed CD19-CAR and representative flow cytometry plots depicting editing outcomes using dsDNA, ssDNA without CTS or with 6-bp tCTS on the 5’ and 3’ ends or 5’ end only. Right side, summary of flow cytometric analysis 4 days after electroporation (n = 7 for TRAC-knock-in into healthy donors). Black lines indicate mean values. Percentage of HDR efficiency and numbers of CAR-expressing cells are shown. For each knock-in condition, 0.5 μg template was used. Statistical analysis was performed using ordinary one-way ANOVA with subsequent Dunn’s correction (for multiple testing) comparing values for each HDRT format with dsDNA as reference. Asterisks represent different p values calculated in the respective statistical test (not significant [ns]: p > 0.5; ∗p < 0.05; ∗∗p < 0.01; ∗∗∗p < 0.001).

**Supplementary Figure 2: Addition of buffer regions and mismatches to the crRNA in the CTS motifs improve CRISPR-Cas12a gene editing outcome with ssDNA donors.** Virus-free insertion of a CD19-CAR transgene into the human CD3ζ locus. (A) Design of ssCTS donor templates is depicted. The insert is flanked by HAs with additional AsCas12a CTS. The CTS comprise a an intact (0-bp mm) or truncated CTS (2- or 4-bp mm), along with a PAM and either no buffer region or a 4-bp buffer region, located on the antisense strand (OS) or on both strands (TS). (B) Left side, schematic of the CD19-CAR receptor. Right side, summary of flow cytometric analyses 4 days after electroporation (n = 6 for CD3ζ-knock-in into healthy donors from 3 independent experiments). Black lines indicate mean values. HDR efficiency and numbers of CAR-expressing cells are shown. For each knock-in condition, 1 μg template was used. Statistical analysis was performed using ordinary one-way ANOVA with subsequent Dunn’s correction (for multiple testing) comparing values for each HDRT format with ssDNA ‘no CTS’ as a reference. Asterisks represent different p values calculated in the respective statistical test (not significant [ns]: p > 0.5; ∗p < 0.05; ∗∗p < 0.01; ∗∗∗p < 0.001).

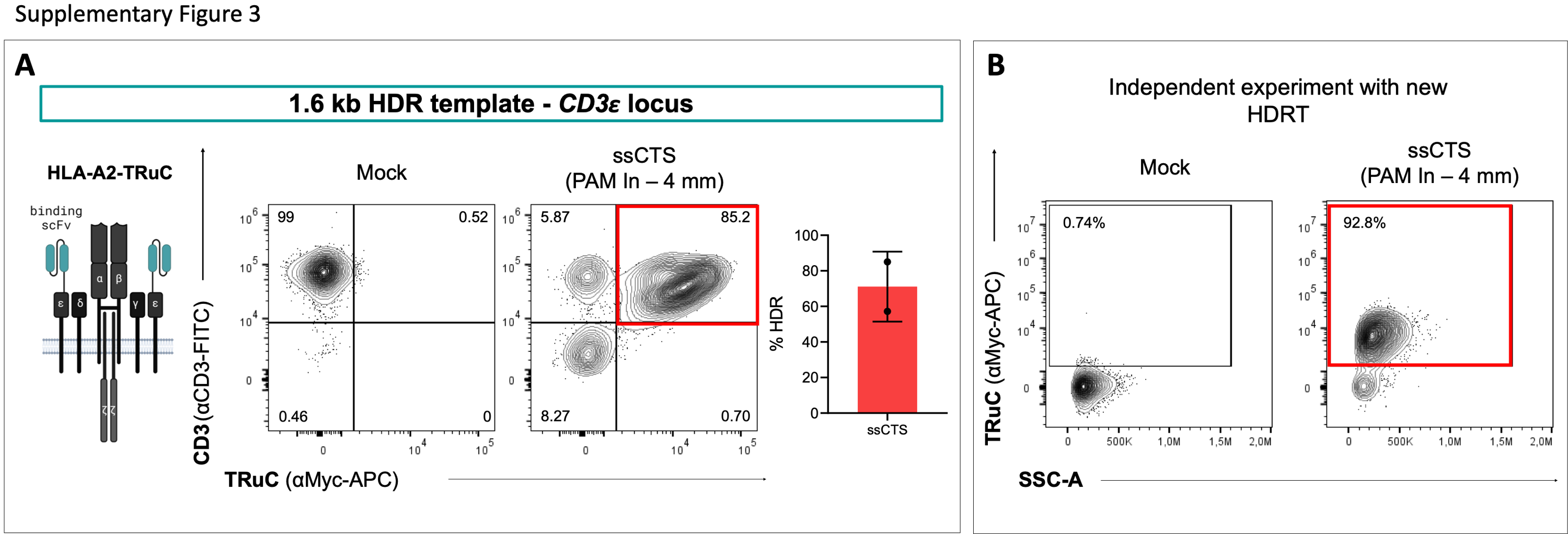

**Supplementary Figure 3: Independent CRISPR-Cas12a knock-in experiments using the HLA-A2-TruC construct confirm the efficacy of the ssCTS templates.** Virus-free insertion of an HLA-A2-TRuC transgene into the human CD3ε locus. (A) Left side, schematic of the expressed receptor and representative flow cytometry plots depicting editing outcomes using the ssCTS HDRT with 4-bp mm from figure 3B. Right side, summary of flow cytometric analysis depicting HDR efficiency 4 days after electroporation (n = 2 for CD3ε -knock-in into healthy donors from one experiment). (B) Independent gene editing experiment using a newly generated ssCTS template batch encoding the same transgene as in A. Flow cytometry plots depicting editing outcomes 4 days after electroporation (n = 1 for CD3ε-knock-in into a healthy donor).
